# A transferrin receptor 1-targeted PNA-peptide conjugate inhibits microRNA-21 expression in cardiac and other mouse tissues

**DOI:** 10.1101/2023.04.13.536802

**Authors:** Genwei Zhang, Sarah Antilla, Chengxi Li, Andrei Loas, Thomas E. Nielsen, Bradley L. Pentelute

## Abstract

MicroRNAs (miRNAs) are implicated in the onset and progression of a variety of diseases. Modulating the expression of specific miRNAs is a possible option for therapeutic intervention. A promising strategy is the use of antisense oligonucleotides (ASOs) to inhibit miRNAs. Targeting ASOs to specific tissues can potentially lower the dosage and improve clinical outcomes by alleviating systemic toxicity. We leverage here automated peptide nucleic acid (PNA) synthesis technology to manufacture an anti-miRNA oligonucleotide (antagomir) covalently attached to a 12-mer peptide that binds to transferrin receptor 1. Our PNA-peptide conjugate is active in cells and animals, effectively inhibiting the expression of miRNA-21 both in cultured mouse cardiomyocytes and different mouse organs (heart, liver, kidney, lung, and spleen), while remaining well-tolerated in animals up to the highest tested dose of 30 mg/kg. Conjugating the targeting ligand to the PNA antagomir significantly improved inhibition of miRNA-21 in the heart by over 50% relative to the PNA alone. Given the modulation of biodistribution observed with our PNA-peptide conjugate, we anticipate this antagomir platform to serve as a starting point for pre-clinical development studies.

**Table of Contents Entry:** 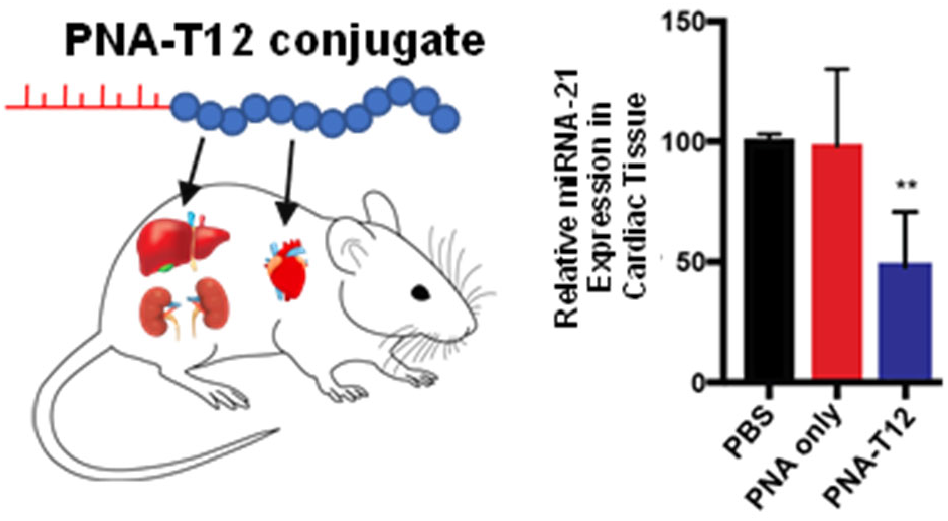

*Synopsis:* Conjugating T12, a peptide targeting transferrin receptor 1 (TfR1), to a peptide nucleic acid (PNA) oligonucleotide targeting microRNA-21 increases delivery of the PNA-T12 conjugate to cardiac tissue relative to PNA alone.

## INTRODUCTION

The implication of many microRNAs (miRNAs) in disease progression has been documented.^1,2^ MiRNA genes are transcribed by RNA polymerase II in the nucleus, producing the precursor miRNAs, termed primary miRNAs, which can be further capped and polyadenylated into pre-miRNA transcripts.^3^ Pre-miRNA typically exists in a hairpin-shaped structure and is processed by the Dicer enzyme into short single-stranded miRNA (mature miRNA) consisting of ∼22 nucleotide bases after being exported into the cytosol via the nuclear pore.^4^ Functional mature miRNA can incorporate into the RNA-induced silencing complex (RISC) in the cytoplasm and complementarily bind the 3’-untranslated region of mRNA targets, regulating protein expression through either mRNA degradation or translational suppression.^3,5^ Over the past two decades, nearly 2,000 precursors and over 2,600 mature miRNAs have been identified and annotated in humans.^6^ Many miRNAs were found to be potent regulators of intracellular signaling pathways and act as important effectors of different cellular functions, such as cell proliferation and differentiation,^7–9^ survival,^10–12^ extracellular matrix production,^13^ and other regulatory events.^1,14–16^ Individual miRNAs can affect the expression of several mRNAs and govern complex biological processes. Not surprisingly, dysregulation of miRNAs was found in some pathophysiological stress and disorder conditions.^11,16–19^

In the diseased state, relevant miRNAs can be either overexpressed or downregulated, therefore presenting opportunities for therapeutic intervention by using either oligonucleotide-based inhibitors or miRNA mimics.^20^ To combat upregulated miRNAs in diseased tissue, targeting strategies with antisense oligonucleotides (ASOs) have gained traction as potential therapies.^12,21,22^ Antagomirs, as a novel class of chemically engineered oligonucleotides, first demonstrated efficacy toward silencing specific miRNAs *in vivo* in 2005^23^ and have shown promise in developing treatments of overexpressed miRNAs.^24–26^ Chemical modifications to the ASO backbone such as phosphorothioate (PS), 2’-*O*-methyl (OMe), 2’-*O*-methoxyethyl (MOE), and locked nucleic acids (LNAs) have greatly improved their metabolic stability, enhanced RNA target specificity, and alleviated off-target toxicity.^27,28^ In other modifications of ASOs, the charged backbone is replaced with neutral components to reduce electrostatic repulsion, such as in phosphorodiamidate morpholino oligomers (PMO) and peptide nucleic acids (PNA). PMOs of at least 15 nucleotides are highly specific, easily soluble, and non-toxic.^27^ Unlike the other ASOs, the backbones of PNAs consist of repeating neutral *N*-(2-aminoethyl)glycine units, rendering them chemically, thermally, and enzymatically stable. Additionally, PNAs exhibit higher binding affinity with their DNA or RNA targets due to their neutral backbone.^27,29^

Recent studies illustrate how delivery of ASOs can safely affect the progression of diseases in humans. Inhibiting miRNA-92a with ASOs led to improved vascularization in myocardial infarction models and more rapid wound healing.^30^ In 2020, the first in-human study of MRG-110, an LNA-DNA mixed antagomir that targets miRNA-92a, showed time- and dose-dependent inhibition of miRNA-92a expression relative to the placebo group after a single dose over the course of twelve weeks.^31^ At the highest dose of 1.5 mg/kg, >95% inhibition of miRNA-92a was observed after 24 h. No significant side effects were reported in the patients participating in this study.^31^ In 2021, the first in-human study was reported of CDR132L, an LNA-based antagomir that targets miRNA-132, which is upregulated in response to cardiac stress, eventually leading to heart failure (HF). Patients in this study received two doses or a placebo four weeks apart. At all doses, rapid and sustained inhibition of miRNA-132 was observed in plasma relative to the placebo group. Importantly, patients receiving at least 1 mg/kg of CDR132L showed plasma levels of miRNA-132 similar to that of a group of healthy volunteers. None of the patients in the study reported adverse side effects.^32,33^ These two studies demonstrate that ASOs are tolerable and effective in humans, augmenting their promise as therapies.

In addition to miRNA-92a and miRNA-132, miRNA-21 is overexpressed in failing hearts, and its inhibition shows promise as a treatment for HF. MiRNA-21 is involved in the extracellular signal-regulated kinase—mitogen-activated protein kinase signaling pathway that modulates apoptotic cell death in cardiac fibroblasts. Increased expression of miRNA-21 reduces the percentage of apoptotic cells of failing hearts, and the greater survival of damaged cells leads to morphology changes exhibited in HF. Silencing miRNA-21 with ASOs in mouse models has shown reduced fibrosis and cardiac dysfunction.^16^ Transverse aortic constriction (TAC) model mice treated with anti-miRNA-21 exhibited higher fractional shortening and decreased cardiac size relative to control mice, partially recovering to healthy measurements.^16^

Despite encouraging results demonstrating the efficacy of ASOs, delivering them to extrahepatic tissues is a challenge because they are often trafficked to the liver or kidney for excretion. Investigating various techniques to modulate the biodistribution of gene therapies is an area of active research. For example, studies have recently shown that *N*-acetylgalactosamine (GalNAc) conjugated to small interfering RNA (siRNA), a double-stranded RNA that activates the RISC complex similarly to miRNAs, can target liver hepatocytes.^20,34^ Targeting the central nervous system and lungs has been achieved by modifying siRNA with lipophilic moieties.^35^ Another strategy leverages the specific binding interactions between antibodies and their target proteins to target non-liver tissue. Antibody-oligonucleotide conjugates (AOCs) were developed that help reduce gene splicing errors in target mRNAs implicated in myotonic dystrophy type 1 (DM1).^36^ Though antibodies are highly specific, they are large (>150 kDa) and complex glycosylated molecules. It has been shown that using only the antigen-binding fragment (Fab) of an antibody to deliver PMOs to muscle tissue can maintain the selectivity of the targeting agent while reducing the molecular weight of the conjugate.^37^ An alternative to antibody-based targeting strategies is employing adeno-associated virus vectors with low pre-existing immunogenicity to target skeletal and cardiac muscle tissue to treat diseases such as Duchenne muscular dystrophy (DMD).^38^ These and other targeting strategies show promising results for improved delivery of gene therapies to desired tissue.

One protein of particular interest for targeted delivery of gene therapies is the transferrin receptor 1 (TfR1). TfR1, or CD71, is highly expressed in cardiac and skeletal muscle tissue and mediates iron transport into cells by binding its native partner transferrin (Tf), which has been used as a metallodrug carrier.^39,40^ Several classes of targeting agents previously discussed are currently being studied in the context of targeting TfR1, including AOCs and Fabs, for the treatment of DMD and other muscular diseases.^36,37^ Centyrins, antigen-specific proteins with specific sites for drug conjugation about 1/15 the size of an antibody, are being investigated as an siRNA delivery agent to treat Pompe disease.^41^ To avoid potentially disrupting iron cycling into cells, there are ongoing efforts to discover bicyclic peptides that target regions of TfR1 that do not bind to Tf.^42^ All of these strategies show significantly improved delivery of the gene therapies to muscle tissue, but they rely on large molecular weight targeting agents that sometimes require high doses for efficacy. For example, one reported Fab complex is effective at 30 mg/kg PMO, corresponding to 68 mg/kg total dosage of compound.^37^ Simplifying the structure of the targeting agent would allow for lower doses to achieve equivalent efficacy, further reducing treatment costs.

Here we report a structurally simple linear 12-mer peptide with nanomolar affinity to TfR1 that alters the biodistribution of anti-miRNA-21 PNA relative to PNA alone. Two peptide binders to TfR1, 12-mer and 7-mer long, respectively, were discovered in 2001 by phage display on whole cells transfected to express TfR1.^43^ Competition assays with Tf indicated that neither binder competes with Tf binding to TfR1. Using phage titer, it was determined that the 12-mer peptide has 15 nM binding affinity to TfR1, while the 7-mer peptide’s binding affinity was only in the micromolar range.^43^ We leverage the strong interaction with TfR1 of this known 12-mer binder, termed T12, to modulate the expression of miRNA-21 *in vitro* and *in vivo* via the delivery of an anti-miRNA-21 PNA. By conjugating T12 to PNA, we observe knockdown of miRNA-21 expression in cardiac tissue *in vivo*, whereas no such activity was observed with unconjugated PNA.

## RESULTS

### Chemically synthesized PNA-T12 antagomir shows selective binding to its pre-miRNA target

Previously, we developed a single-shot strategy to synthesize peptide-conjugated PNAs on an automated platform,^44^ which was controlled using a modular script in the ‘Mechwolf’ programming environment (Figure1a).^26^ The PNA sequence was reverse-complemented to specifically target miRNA-21. Synthetic T12, amino acid sequence THRPPMWSPVWP,^43^ was verified to exhibit nanomolar binding affinity to the recombinant TfR1 ectodomain (residues C89-F763) by bio-layer interferometry (BLI, Figure 1b). We observed an apparent dissociation constant *K*_D_ = 26 nM, in line with the previously reported value of 15 nM determined by phage titer.^43^ For the PNA-T12 conjugate, we included a simple Lys_3_ linker between the two components, resulting in the final sequence of CATCAGTCTGATAAGCTA-KKK-THRPPMWSPVWP-CONH_2_. After multiple rounds of synthesis and purification, >10 mg of PNA-T12 (>95% purity by LC-MS) were isolated. Following sample preparation, a gel shift assay was performed to demonstrate that chemically synthesized PNA-T12 could selectively bind to the target miRNA sequence. As shown in Figure 1c, the anti-miRNA-21 PNA-T12 construct bound and shifted pre-miRNA-21, but not pre-miRNA-34a used as a control, indicating miRNA target selectivity.

**Figure 1.**
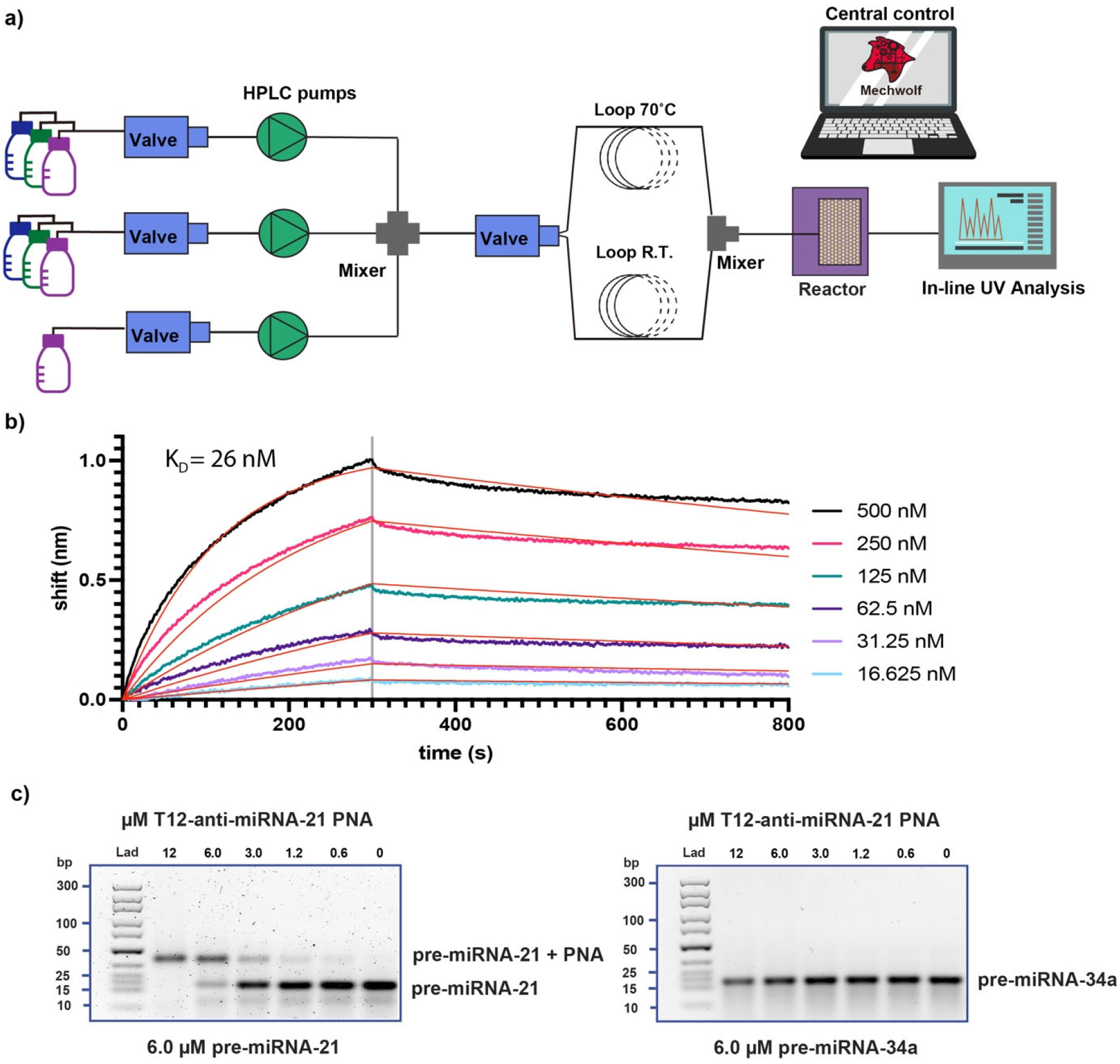
Chemically synthesized PNA-T12 conjugate binds the targeted pre-miRNA. (a) T12-conjugated PNA was prepared by automated single-shot flow synthesis. (b) Synthetic T12 exhibits concentration-dependent binding to recombinant TfR1 ectodomain with nanomolar affinity by BLI. Biotinylated T12 loaded on streptavidin (SA) tips at 1.5 µM was allowed to associate with TfR1 at the above concentrations for 300 s, followed by a dissociation step for 500 s. (c) Gel shift assays show the selective binding between chemically synthesized anti-miRNA-21 PNA-T12 and pre-miRNA-21, with pre-miRNA-34a used as a control. For each gel, 6 µM miRNA was pre-mixed with 0-12 µM PNA-T12 at 37°C for 1 hour and loaded into a 5% agarose gel at 400 ng miRNA/well. Gels were run at 100 V for 70 min in 1X TBE buffer.

### Anti-miRNA-21 PNA-T12 conjugate inhibits miRNA-21 expression in mouse cardiomyocytes

To assess the efficacy of our PNA-T12 conjugate for gene silencing, we performed a miRNA inhibition efficacy assay in a cardiac muscle cell line (HL-1). HL-1 cells were derived from the AT-1 mouse atrial cardiomyocyte tumor lineage and can be repeatedly passaged while maintaining a cardiac-specific phenotype.^45^ Stable HL-1 cells were incubated with PNA-T12 or PNA alone at gradually increasing concentrations (for PNA alone 0.01, 0.1, 1, and 10 µM; for PNA-T12 0.1, 0.5, 1, and 5 µM). One day later, the attached cells were scraped off the plate and harvested for RNA extraction, followed by reverse transcription and quantitative real-time polymerase chain reaction (qRT-PCR). The mouse miRNA-16 was selected as an internal control for the qRT-PCR reactions based on a previous report,^46^ and the comparative C-T method was used to calculate the cycle threshold.^47^ A clear dose-dependent reduction in miRNA-21 expression, up to >50% inhibition at 5µM, was observed in the cells incubated with PNA-T12 (Figure 2b). However, no dose-dependent inhibition was observed in the cells incubated with PNA alone (Figure 2a). These results indicate that conjugating the targeting ligand T12 to PNA improves uptake and intracellular delivery of PNA in cardiac cells and drives targeted miRNA silencing.

**Figure 2.**
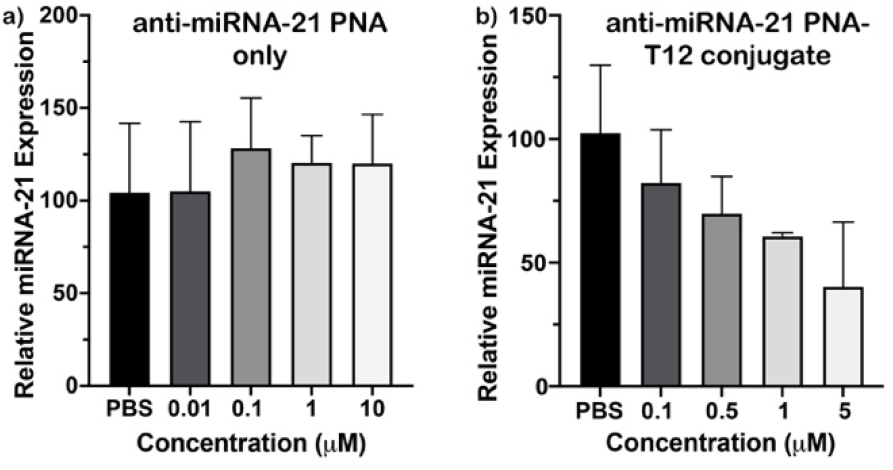
T12-conjugated PNA antagomir displays miRNA-21 inhibition in mouse cardiomyocytes. The bioactivity of the synthetic PNA-T12 conjugate was tested in cultured HL-1 mouse cardiomyocytes, and the targeted miRNA inhibition efficacy was measured via qRT-PCR. HL-1 cells were incubated at 37°C in complete Claycomb media for 24-36 h after being dosed with PNA or PBS. Anti-miRNA-21 PNA conjugated to T12 shows dose-dependent inhibition of miRNA-21 relative to the PBS control when conjugated to T12 (b), but not on its own (a). For each condition, N=3 technical replicates were used, with relative expression calculated using the comparative *C*_*T*_ method. Statistical significance is calculated using Student’s t-test, ** p < 0.01; **** p < 0.0001; n.s. not significant.

### The anti-miRNA-21 PNA-T12 conjugate affects miRNA-21 inhibition in the mouse heart

Minimizing off-target delivery could improve the therapeutic window of antagomir PNAs. We investigated the PNA targeting capabilities of the T12 ligand *in vivo* following the improved uptake in HL-1 mouse cardiomyocytes observed *in vitro*. Two experimental groups of wild-type mice (strain C57BL/6, n = 5) were dosed with 30 mg/kg or 15 mg/kg of PNA-T12 via retro-orbital injection. Two control groups (n = 5) were injected with equivalent volume of 1X phosphate-buffered saline (PBS) and 30 mg/kg PNA alone. All mice were sacrificed two weeks after compound administration. At the time of sacrifice, all mice appeared healthy, and we observed no significant adverse side effects in the physical appearance or behavior of the mice. To measure the efficacy of PNA-T12 toward miRNA-21 inhibition, five organs (heart, liver, kidney, lung, and spleen, Figure 3a) were extracted from dissected mice and preserved in Trizol reagent for subsequent analysis. Total RNA was extracted from the preserved organ samples and qRT-PCR was performed on each sample using primers provided within the Taqman kit. We observed the naked PNA construct at 30 mg/kg caused significant miRNA-21 reduction in the kidney and liver, but not in the heart (Figure 3b-d), which reinforces the long-standing challenge of delivering ASO molecules outside of vascularized tissues.^48,49^ In contrast, PNA-T12 inhibits approximately 50% of miRNA-21 expression in cardiac tissue at 30 mg/kg (Figure 3b). In lung tissue, including a targeting ligand reduces inhibition of miRNA-21 expression relative to PNA alone (Figure 3e), indicating that T12 is able to alter the biodistribution of PNA in mice. At 15 mg/kg, there is reduced gene silencing in cardiac liver, kidney, and spleen tissue (Figure 3b,c,d,f), consistent with dose dependence. While this outcome is promising in demonstrating that lower doses of these compounds can reduce off-target delivery, no significant efficacy was observed in the cardiac tissue below 30 mg/kg. Nonetheless, at 30 mg/kg we were able to enhance miRNA-21inhibition in the cardiac tissue of mice using a directed targeting strategy without eliciting observable side effects. This indicates that at the doses studied, our PNA-T12 conjugate is effective and well-tolerated in mice. Our peptide-conjugated antisense PNA construct opens a route to produce lead compounds to continue in-depth pre-clinical investigations.

**Figure 3.**
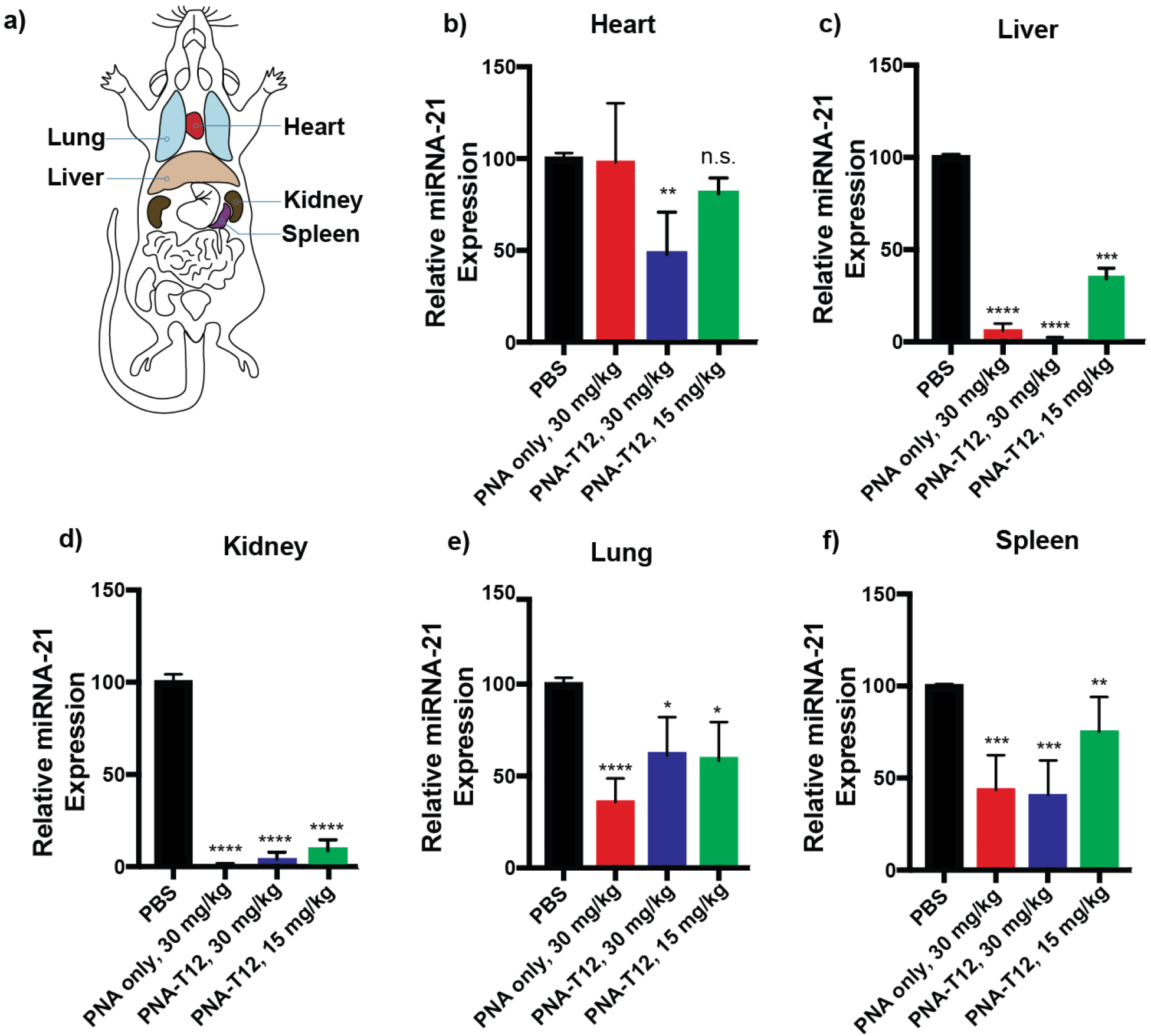
T12-conjugated anti-miRNA-21 PNA exhibits altered biodistribution relative to PNA alone. (a) Schematic representation of the mouse organs dissected to assess miRNA-21 inhibition. Anti-miRNA-21 PNA alone drives substantial inhibition of miRNA-21 in the mouse liver (c) and kidney (d), but not in the heart (b). In contrast, T12-conjugated anti-miRNA-21 PNA renders reduced expression of miRNA-21 at equivalent doses in mouse cardiac tissue (b) along with a diminished inhibitory effect in lung tissue (e). The efficacy of the PNA-T12 conjugate in mouse cardiac tissue is reduced at doses below 30 mg/kg (b). Statistical significance is calculated using Student’s t-test, ** p < 0.01; **** p < 0.0001; n.s. not significant.

## DISCUSSION

In this study we show that a TfR1-targeting ligand can modulate the biodistribution of ASOs *in vivo* in a beneficial manner. Addressing the root biological causes of several diseases that are driven by upregulation or downregulation of RNA expression could lead to reversal of the disease progression rather than managing symptoms. However, because of the high concentration of receptors that facilitate the uptake of ASOs in vascularized tissue such as liver, kidney, and spleen, targeted delivery of ASO-based therapeutics remains a challenge.^48^ This proof-of-concept study illustrates that a simple receptor-targeting peptide has the capacity to modulate the activity of a conjugated ASO cargo toward miRNA expression in cardiac tissue.

This small linear peptide performs on a comparable level with much larger, more complex targeting agents. The size and structural simplicity of T12 is reliably accessed with flow synthesis in hours,^44^ offering a more rapid and straightforward production method relative to recombinant expression and other bioengineering procedures required for acquiring antibodies and other complex macromolecular targeting agents. Protein and antibody-based targeting agents typically require doses in the range of 1-30 mg/kg for efficacy.^36,37,41^ The best reported efficacy for a Fab-PMO conjugate is achieved at 30 mg/kg of ASO equivalent, which corresponds to nearly 70 mg/kg of total compound,^37^ a large amount of material required for human patients. Our PNA-T12 conjugate demonstrated about 50% efficacy in cardiac tissue at 30 mg/kg total compound. Thus, T12 alters the biodistribution of ASOs at a competitive level with other larger targeting modalities. Looking ahead, we seek to further study the interaction between T12 and TfR1 to reduce the dose of conjugates required for efficacy.

Current advances in the field of ASO targeting are expanding the viability of RNA interference (RNAi) therapeutics. As the number of target miRNA sequences in the progression of various diseases grows, ASOs and other gene therapies are becoming a key area of interest.^11,16–19^ Several ongoing efforts as well as this work demonstrate that active targeting can be used to beneficially direct where ASOs or other RNAi therapeutics are delivered in the body.^36–38,41,42^ The development of rapid flow synthesis of peptides, ASOs, and their conjugates will allow for straightforward manufacturing of many candidate compounds on a short time scale not achievable by biological methods for larger, more complex conjugates.^26,44^ These efforts have shown encouraging progress in achieving extrahepatic targeted delivery of gene therapies. As targeting strategies improve and off-target delivery is reduced, ASOs can become much more prevalent and widely available treatments for diseases.

## SUPPORTING INFORMATION

Experimental procedures for LC-MS traces, gel shift assays, BLI assays, and qRT-PCR efficacy assays, and characterization data for all compounds. The Supporting Information is available free of charge on the ACS Publications website at DOI: 10.1021/xxx.

## NOTES

The authors declare the following competing financial interests: B.L.P. is a co-founder and/or member of the scientific advisory board of several companies focusing on the development of protein and peptide therapeutics. The authors are preparing a provisional patent disclosure regarding the methodology and compounds described in this study. The Python code for automated operation of the flow synthesis instrument is available in a GitHub repository (https://github.com/L-Chengxi/MechWolf_Pull).

## ACKNOWLEDGEMENTS

Financial support for this work was provided by Novo Nordisk A/S (to B.L.P.). We acknowledge the Preclinical Modeling, Imaging and Testing Facility at the MIT Koch Institute’s Robert A. Swanson (1969) Biotechnology Center for technical support in the handling and care of animals (NCI Core Center Grant P30-CA14051). We thank the Division of Comparative Medicine at MIT and in particular Sarah Elmiligy and Virginia Spanoudaki for their assistance with animal studies and training; Joseph S. Brown and Michael A. Lee for insightful discussions and suggestions; and Andrew Wilson for his help in the design and production of the automated flow instrument.

