## Supporting Information for "A transferrin receptor 1-targeted PNA-peptide conjugate inhibits microRNA-21 expression in cardiac and other mouse tissues"

### 1. General information

Throughout this work, no unexpected or unusually high safety hazards were encountered.

*Synthesis reagents*

H-Rink Amide (0.50 mmol/g loading or 0.18 mmol/g) resin for peptide and PNA synthesis were purchased from PCAS Biomatrix. All the fluorenylmethyloxycarbonyl (Fmoc) protected amino acids were purchased from Novabiochem-line from Sigma Millipore and used as received. All the Fmoc-protected PNA monomers (moA, moC, moG, and moT) were purchased from PNA bio; O-(7-azabenzotriazol-1-yl)-*N*,*N*,*N*',*N*'-tetramethyluronium hexafluorophosphate (HATU, ≥97.0%) was purchased from P3 Biosystems; *N,N,N',N'*-tetramethyl-O-(1H-benzotriazol-1-yl)uronium hexafluoro-phosphate (HBTU, ≥97.0%) was purchased from P3 Biosystems; Diisopropylethylamine (DIEA; 99.5%, biotech grade, catalog number 387649), piperidine (≥99.0%), piperazine (≥99.0%), morpholine (≥99.5%), trifluoroacetic acid (HPLC grade, ≥99.0%), triisopropylsilane (≥98.0%), formic acid (FA, ≥95.0%), and 1,2-ethanedithiol (EDT, GC grade, ≥98.0%) were purchased from Sigma-Aldrich. Diisopropylethylamine (DIEA; 99.5%, biotech grade, catalog number 387649), piperidine (≥99.0%) and formic acid (FA, ≥95.0%) were purchased from Sigma-Aldrich. *N*, *N*-dimethylformamide (DMF, Biosynthesis OmniSolv® grade) was purchased from EMD Millipore (DX1732-1); AldraAmine trapping agents (for 1000~4000 mL DMF, catalog number Z511706) were purchased from Sigma-Aldrich.

*Cleavage reagents*

Trifluoroacetic acid (TFA, HPLC grade, ≥99.0%), triisopropylsilane (TIPS, ≥98.0%), and 1,2-ethanedithiol (EDT, GC grade, ≥98.0%) were purchased from Sigma-Aldrich.

*Analysis and purification reagents*

Water for HPLC was purified to 18.2MΩ/cm resistivity on a Millipore Milli-Q system. Acetonitrile (HPLC-grade) was purchased from VWR International (Philadelphia, PA) and acetonitrile (LC-MS grade) was purchased from Sigma-Aldrich (St. Louis, MO).

*Cell assay reagents*

Claycomb medium, (+/-)-norepinephrine (+)-bitartrate salt, L-ascorbic acid, L-glutamine solution bioxtra, trypsin-EDTA solution, trypsin inhibitor, fibronectin from bovine plasma, and gelatin from bovine skin type B were all purchased from Sigma-Aldrich; fetal bovine serum was purchased from Thermo Fisher Scientific.

*Animal experiments*

Wild-type mice (strain C57BL/6) were ordered from Charles River; Dulbecco's phosphate buffered saline was purchased from Sigma-Aldrich; TRIzol reagent, TaqMan universal PCR master mix, and TaqMan MicroRNA assay kits were all purchased from Invitrogen; Anti-MiR21 LNA molecule was ordered from Integrated DNA Technologies.

### 2. Procedure for automated PNA/PNA-peptide synthesis

Unmodified PNAs and peptide-conjugated PNAs were synthesized on an automated synthesizer named Tiny Tides[1] following our reported protocols. [2] We redescribe the procedure below for the reader:

*Unmodified PNA synthesis*

The automated synthesis of PNAs was performed using the self-designed oligonucleotide synthesizer (Tiny Tides). Rink amide resin (15 mg, 0.5 mmol/g loading) was loaded into the reactor. The reactor was connected to the reactor head and heated to 70 ˚C. DMF was delivered at 5 mL/min (2.5 mL/min per pump) for 20 seconds to remove air. The flow was stopped and the resin was allowed to swell at 70 ˚C for 5 minutes. The flow protocol was started with an initial DMF wash at 5 mL/min (2.5 mL/min per pump) 70 ˚C for 40 seconds, then coupling solution composed of one-part 0.2 M PNA monomer subunit in DMF, one-part 0.18 M HBTU in DMF, and one-part 10% DIEA (v/v) in DMF was delivered for 10 seconds (10 eq PNA monomer) at 70 ˚C. Next, DMF was delivered at 5 mL/min (2.5 mL/min per pump) under 70 ˚C for 20 seconds to ensure all the monomer solutions arrived at the reactor and clean the loop. The 6-position valve then switched to the room-temperature loop. DMF was delivered at 5 mL/min (2.5 mL/min per pump) for 40 seconds at room-temperature. Cold DMF flow (rt) mixed with the hot reactor (70 ˚C) generated an in-situ 40 ˚C environment for deprotection. Deprotection was performed with one-part 40% piperidine, 2% formic acid (v/v) in DMF, and one-part DMF for 50 seconds in the room-temperature loop. After a 20-second room temperature DMF wash, the 6-position valve was switched to the 70 ˚C loop. DMF was delivered at 5 mL/min for 40 seconds to wash the resin and preheat the reactor. No capping or multiple couplings were needed, and each single coupling cycle took 3 minutes. Repeat the synthesis cycle to finish the long PNA chain assembly.

*PNA-T12 synthesis*

For single-shot T12 conjugated PNA synthesis, T12 was pre-synthesized on the H-Rink Amide resin (0.5 mmol/g) with a peptide synthesizer developed in our lab.[3-5] The T12 bound resin was used directly for automated PNA-T12 synthesis on Tiny Tides. Both $(Lys)_3$ linker and PNA sequences were synthesized directly on Tiny Tides.

### 3. Cleavage protocols

After synthesis, the PNA bound resin was washed with dichloromethane (3 x 5 mL), dried in a vacuum chamber, and weighed. Then the resin was transferred into a 15 mL conical polypropylene tube. Approximately 1 mL of cleavage solution (94% TFA, 1% TIPS, 2.5% EDT, 2.5% water) was added to the tube and kept at room temperature for 2 hours. After cleavage, the resin was removed by filtration, the filtrate was concentrated under a stream of nitrogen and the PNA product was precipitated in dry ice-cold diethyl ether (12 mL) by centrifugation and washed three times. The supernatant was discarded, and the residual was dissolved in 50% acetonitrile in water with 0.1% TFA. The PNA solution was filtered with a Nylon 0.22 µm syringe filter, frozen with liquid nitrogen, and lyophilized to dried powder. Finally, the crude PNA was weighed.[2]

### 4. Purification protocols

***Method A:***

HPLC purification was carried out on a reversed-phase preparative HPLC using an Agilent mass directed purification system (1260 Infinity LC and 6130 single quad MS) with UV detection at 260 nm. Column: Agilent Zorbax SB-C3 (9.4 x 250 mm, 5 µm). Flow rate 4.0 mL/min; Temperature: 60 ˚C. Solvent System: A linear gradient of acetonitrile with a 0.1% TFA additive (solvent B) in water with a 0.1% TFA additive (solvent A) was used. Gradient: 5 min hold 1% B, 1-31% B gradient from 5 to 105 min, 31-65% B gradient from 105 to 120 min, hold 65% B from 120 to 125 min. A final 5.5 min hold was performed with 1% B. The method in total lasted 125 min. Fractions were collected every minute.

***Method B:***

Reverse phase purification was carried out on a Biotage Selekt flash purification system with UV detection at 280 and 214 nm. Column: Biotage Sfär C18 (12g column size, particle size 20 µM, pore size 300 Å, column volume (CV) 17 mL); Flow rate 12 mL/min; Temperature: r.t. Solvent System: A linear gradient of acetonitrile with a 0.1% TFA additive (solvent B) in water with a 0.1% TFA additive (solvent A) was used. Gradient: 1 CV hold 5% B, 5-15% B gradient over 1 CV, 15-45% B gradient over 10 CV, 45-95% B gradient over 0.5 CV, hold 95% B for 2 CV, return to 5% B over 0.1 CV, and hold 5% B for 1 CV. Fractions were collected by UV peak detection, with thresholds of 15 mAu at 280 nm and 50 mAu at 214 nm.[2]

### 5. Liquid chromatography-mass spectrometry (LC-MS) analysis

Analysis was performed on an Agilent 1290 Infinity HPLC coupled to an Agilent 6550 Q-TOF with Dual Jet Stream ESI ionization and iFunnel. MS was run in positive ionization mode, extended dynamic range (2 GHz), and low mass range (m/z in range 100 to 1700). All peptide and PNA solutions were filtered and then diluted to approximately 0.1 mg/mL before sample loading. Buffer A: 0.1% formic acid in $H_2O$. Buffer B: 0.1% formic acid in acetonitrile.

The following LC-MS methods were used at different situation:

***Method A:***

Column: Phenomenex Aeris C4 (2) (3.6 um, 2.1 x 150 mm, 100 Å silica); Flow Rate: 0.2 mL/min; Gradient: 1% B 0-1.50 min, linearly ramp from 1% B to 61% B 1.50 to 6.50 min, hold 90% B from 6.51 to 8 min. Post time is 1% B for 5 min.

***Method B:***

Column: Phenomenex Aeris C4 (2) (3.6 um, 2.1 x 150 mm, 100 Å silica); Flow Rate: 0.2 mL/min; Gradient: 1% B 0-2 min, linearly ramp from 1% B to 61% B 1.50 to 10 min, hold 90% B from 10.1 to 12 min. Post time is 1% B for 5 min.

***Method C:***

Column: Agilent ZORBAX 300SB C3 (5 um, 2.1 x 150 mm); Flow Rate: 0.5 mL/min; Gradient: 1% B 0-2 min, linearly ramp from 1% B to 91% B 2 to 12 min. Post time is 1% B for 3 min.

### 6. Biotinylated cardiac targeting ligand binding affinity study

Peptide binding validation was carried out using bio-layer interferometry (BLI) on a Gator Bio GatorPlus system. The plate agitation speed was kept at 1,000 rpm and the temperature was maintained constantly at 30 °C. Streptavidin (SA)-coated biosensor tips were used to load N-terminal biotinylated cardiac targeting ligand (CTL) T12 (sequence: THRPPMWSPVWP) which was dissolved at 1.5 μM in a kinetic buffer (K.B.): 1 × PBS with 0.1% BSA and 0.02% tween20. After loading the peptide for 120 s, the biosensor tips were then moved into solutions containing various concentrations (15.625 nM, 31.25 nM, 62.5 nM, 125 nM, 250 nM, and 500 nM) of recombinant transferrin receptor (TfR1/CD71) protein (purchased from Sino Biological) in the K.B. to obtain the association curve. After a 300-s association step, the tips were moved back into the K.B. for 500 s to collect the dissociation curve. Peptide-only (1.5 μM) and protein-only (500 nM) conditions were used as references for background subtraction. The association and dissociation curves were analyzed in the Gator Bio GatorPlus Data Analysis Software with defined parameters (linked global kinetic fitting algorithm, binding model 1:1) to calculate the apparent dissociation constant ($K_D$). The analysis results were shown in Figure 1, the $K_D$ value is 26 nM (reported $K_D$ = 15 nM by Scatchard plot from phage titer).[6]

### 7. Gel shift assay

Synthetic purified PNA-T12 was pre-mixed with 6 μM pre-miRNAA-21 or pre-miRNA-34a (ordered from IDT) at different concentrations (0, 0.6, 1.2, 3.0, 6.0 and 12 μM) for 1 hour at 37 °C. Subsequently, a 5% agarose gel was used for gel band separation and analysis, the gel running condition was 100 V for 70 min in 1×TBE buffer. The gel was imaged with SYBR gold nucleic acid stain purchased from Thermo Fisher.

### 8. Bioactivity study of Anti-MiRNA PNAs in HL-1 cells

A cardiac muscle cell line derived from AT-1 mouse atrial cardiomyocyte tumor lineage, named HL-1, was cultured at 37 °C and 5% $CO_2$ in complete Claycomb medium until 80% confluency. Purified PNA-T12 constructs with serial concentrations were added into adherent HL-1 cells. After 24-36 hours inoculation, the total RNA was extracted from treated cells using TRIzol reagent following manufacture's instruction (also specified below). The TaqMan microRNA assay kit was used for quantitative real-time polymerase chain reaction (qRT-PCR). After data acquisition and cycle threshold calculation, the expression levels of relevant miRNAs were calculated using a comparative $C_T$ method (in comparison with a PBS-treated control after being internally normalized to a housing keeping microRNA, MiR-16).

*Extract RNA*

Add 500 μL TRIzol Reagent to cell pellets, then incubate for 5 min at room temperature to allow complete disruption of the cell membrane. Add 200 μL of chloroform per 1 mL of TRIzol reagent used for lysis, then securely cap the tube, and thoroughly mix by shaking. Incubate for 3 minutes. Centrifuge the sample for 15 minutes at 12,000 g at 4 ˚C. Transfer the upper colorless aqueous phase containing the RNA to a new tube carefully with a micropipette.

*Isolate RNA*

Add 500 μL of isopropanol to the aqueous phase, incubate for 10 min at 4 ˚C. Centrifuge the solution for 10 minutes at 12,000 g at 4 ˚C. Note: the total RNA precipitate forms a white gel-like pellet at the bottom of the tube. Discard the supernatant with a micropipette.

*Wash RNA*

Resuspend the pellet in 1 mL of 75% ethanol, and vortex the sample briefly. Then, centrifuge the sample for 5 minutes at 7500 g at 4 ˚C. Discard the supernatant with a micropipette. Air dry the RNA pellet for 10-30 minutes. Note: Do not let the pellet very dry to make sure the solubility of RNA.

*Reverse transcription (RT)*

TaqMan™ MicroRNA Reverse Transcription Kit was used for reverse transcription. The concentration of each RNA sample was measured using a BioTek Epoch microplate spectrophotometer. Prepare 50 ng/μL RNA solution according to the UV concentration. To a 96-well plate, add 0.1 μL dNTP substrate (100 mM wdTTP), 0.5 μL reverse transcriptase, 1 μL RT buffer (10X), 0.14 μL RNase inhibitor (20 U/μL), 5.25 μL nuclease-free water, 2 μL primer, and 1 μL RNA sample to each well. MiRNA-16 was used as an internal control. Perform the reverse transcription on a regular PCR cycler as follows: 16 °C for 30mins, 42 °C for 30mins, 85 °C for 5 min, and then chilled at 4 °C until further manipulation.

*qRT-PCR*

The qRT-PCR experiment was performed using the TaqMan microRNA assay kit and the PCR 2X super mix solution with three technical repeats on a Roche light cycler 480 instrument at MIT Proteomics Core Facility. The qRT-PCR running condition is: 95˚C – 10s, 95˚C – 15s, 60˚C – 60s, and repeat for 50 cycles. The fold change relative to control was calculated using the comparative $C_T$ method, and equation was specified as follows:[7]

$$\text{Fold change} = 2^{-[\Delta CP sample - \Delta CP control]}$$

### 9. T12-conjugated PNA efficacy study in mice

Wild-type mice were reared under a 12-hour light and 12-hour dark cycle and housed at the facility within MIT Division of Comparative Medicine (DCM). To study the PNA-T12 *in vivo* efficacy, PNA-T12 was administered into wild-type mice via retro-orbital injection. Two experimental groups of mice (n=5) were treated with 30 mg/kg and 15 mg/kg PNA-T12, and two control groups of mice (n=5) were injected with equivalent volume of 1× PBS and 30 mg/kg PNA only. At day 14 post-injection, the mice were euthanized and sacrificed using approved protocols and in compliance with the MIT Committee on Animal Care (CAC) guidelines. To measure the efficacy of PNA-T12 injections, five mouse organs (heart, liver, kidney, liver, and spleen) from the mice were dissected, collected, and preserved in TRIzol Reagent for subsequent analysis.

Homogenize the organs in TRIzol using a Dounce homogenizer, incubate for 5 minutes to allow complete dissociation of the nucleoproteins complex. RNA extraction, reverse transcription and qRT-PCR were performed similarly following the procedures outlined in Section 8.

### 11. LC-MS data of synthesized PNA samples

All the PNA samples were synthesized and purified according to the protocols in Supporting Information Section 2 and analyzed with LC-MS before use.

***Figure S1.*** Sample: Anti-MiRNA-21 PNA
Sequence: CATCAGTCTGATAAGCTA-KKK-$CONH_2$.
Synthesis method: Automated flow synthesis.
Purification method: Section S4, method A
LC-MS method: Section S5, method A
Calculated: 5263.27 Da
Observed: 5263.55 Da

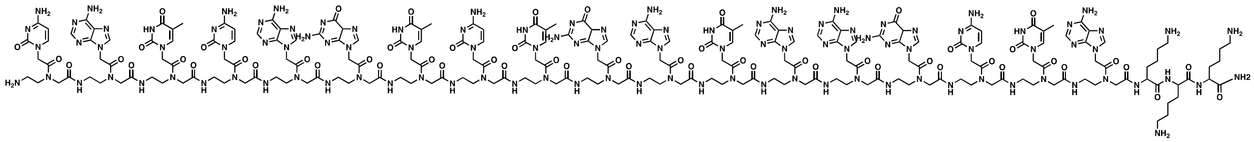

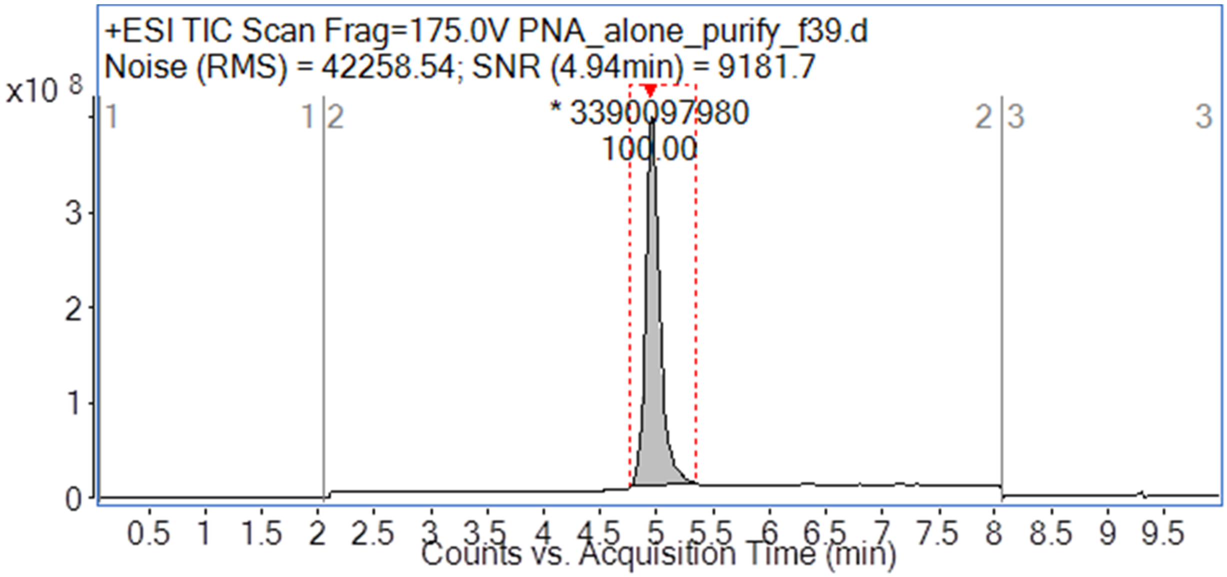

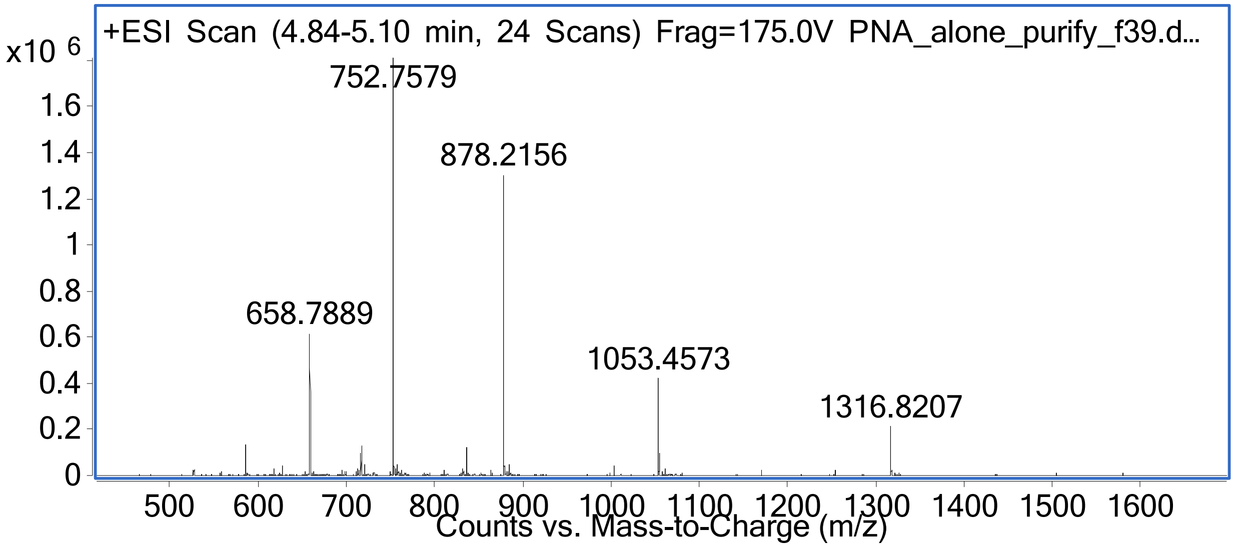
+ESI Scan (4.84-5.10 min, 24 Scans) Frag=175.0V PNA_alone_purify_f39.d...
x10 6
1.6
1.4
1.2
1
0.8
0.6
0.4
0.2
0
752.7579
878.2156
658.7889
1053.4573
1316.8207
500 600 700 800 900 1000 1100 1200 1300 1400 1500 1600
Counts vs. Mass-to-Charge (m/z)

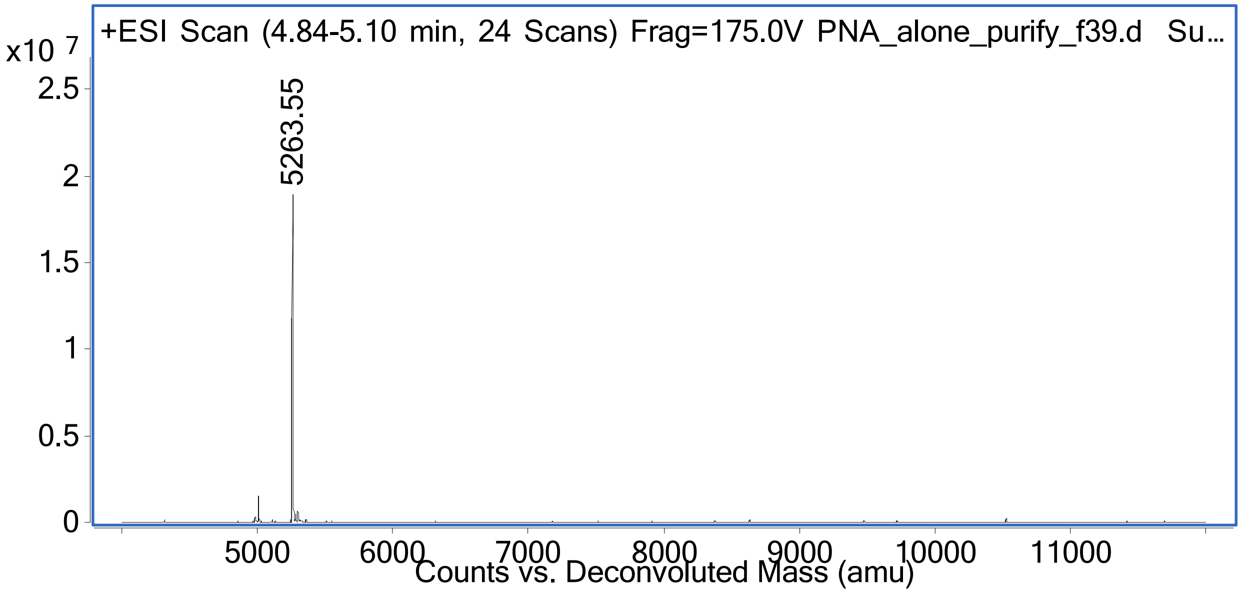
+ESI Scan (4.84-5.10 min, 24 Scans) Frag=175.0V PNA_alone_purify_f39.d Su...
x10 7
2.5
2
1.5
1
0.5
0
5263.55
5000 6000 7000 8000 9000 10000 11000
Counts vs. Deconvoluted Mass (amu)

***Figure S2.*** Sample: Anti-MiRNA-21-T12
Sequence: CATCAGTCTGATAAGCTA-KKK-THRPPMWSPVWP-$CONH_2$.
Synthesis method: Automated flow synthesis.
Purification method: Section 4, method A
LC-MS method: Section S5, method B
Calculated: 6736.01 Da
Observed: 6736.07 Da

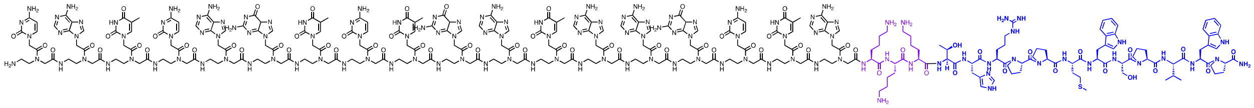

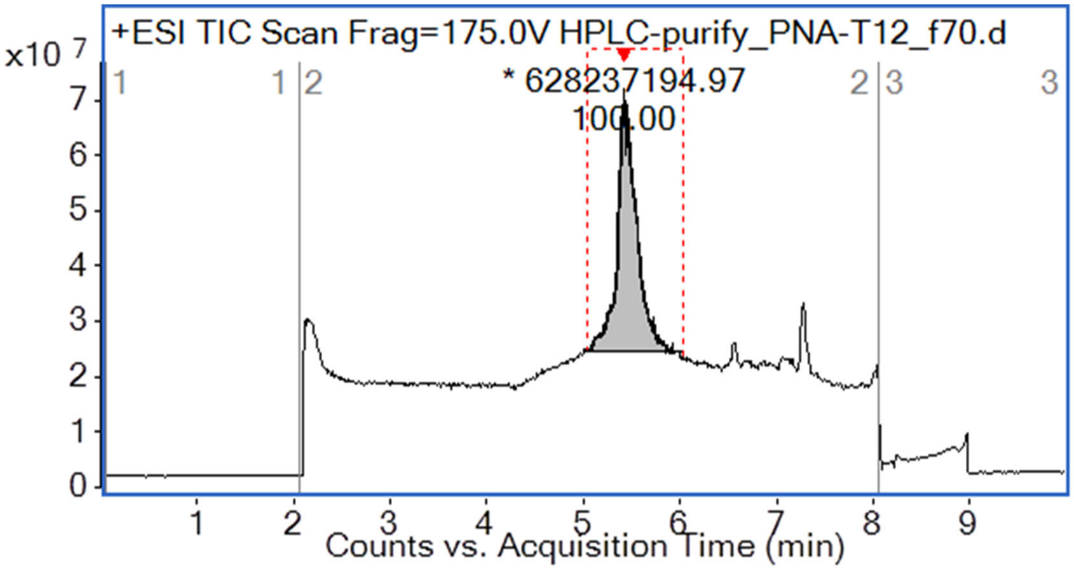

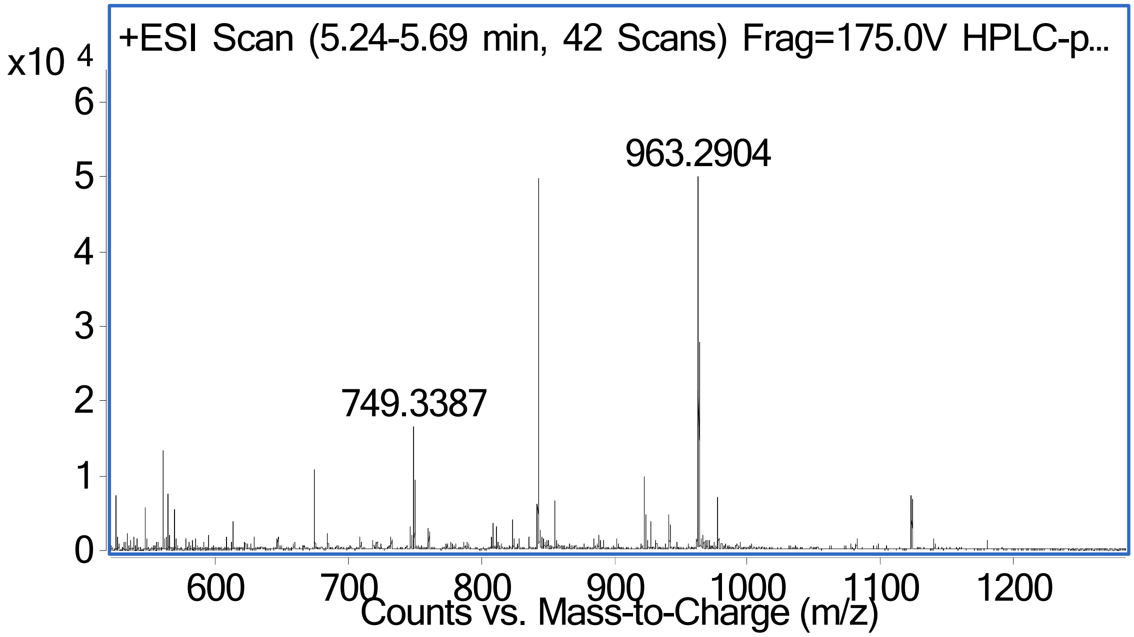

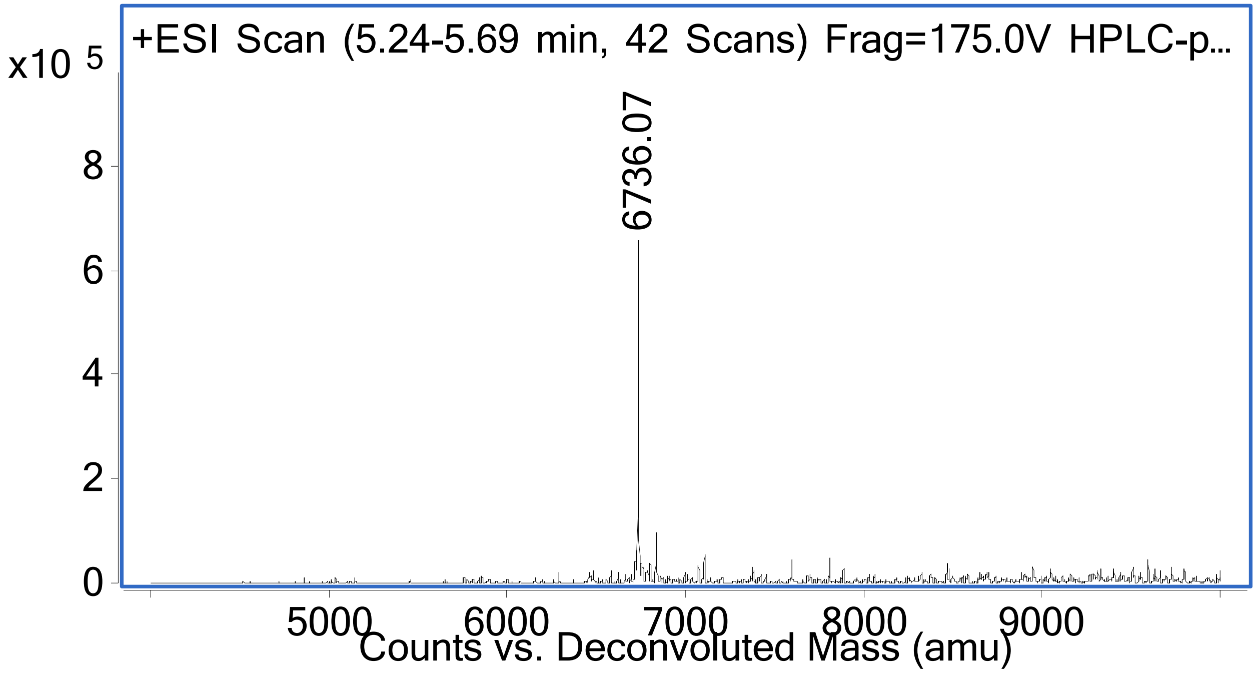
+ESI Scan (5.24-5.69 min, 42 Scans) Frag=175.0V HPLC-p...
x10 5
8
6
4
2
0
6736.07
5000
6000
7000
8000
9000
Counts vs. Deconvoluted Mass (amu)

***Figure S3.*** Sample: biotin-T12

Sequence: biotin-PEG4-THRPPMWSPVWP-CONH2.

Synthesis method: Automated flow synthesis.

Purification method: Section 4, method B

LC-MS method: Section S5, method C

Calculated: 1963.35 Da

Observed: 1962.95 Da

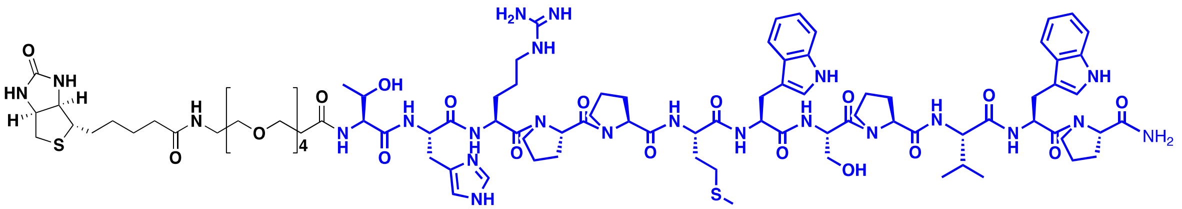

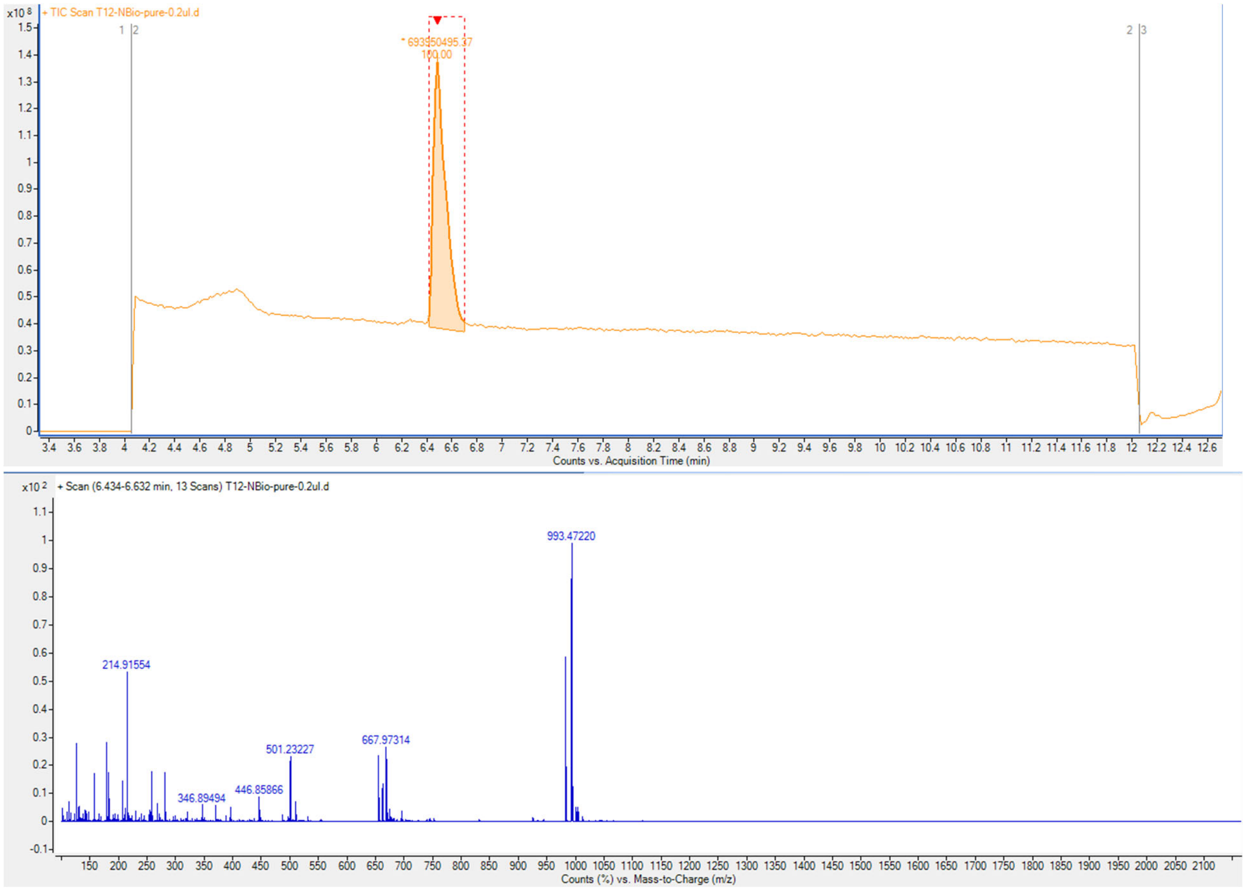
